# The kinesin-like protein Pavarotti functions non-canonically to regulate actin dynamics

**DOI:** 10.1101/561936

**Authors:** Mitsutoshi Nakamura, Jeffrey M. Verboon, Clara L. Prentiss, Susan M. Parkhurst

**Affiliations:** Basic Sciences Division, Fred Hutchinson Cancer Research Center, Seattle, WA, USA 98109

**Keywords:** Pavarotti, Tumbleweed, Pebble, RhoGAP, RhoGEF, actin, wound repair, oogenesis, cytokinesis, Drosophila

## Abstract

Pavarotti, the Drosophila MKLP1 ortholog, is a kinesin-like protein that with Tumbleweed (MgcRacGAP) works together as the centralspindlin complex. This complex is essential for cytokinesis where it helps to organize the contractile actomyosin ring at the equator of dividing cells by activating the RhoGEF Pebble. Actomyosin rings also function as the driving force during cell wound repair. We previously showed that Tumbleweed and Pebble are required for the cell wound repair process. Here, we show that Pavarotti also functions during wound repair and confirm that while Pavarotti, Tumbleweed, and Pebble are utilized during this cellular repair, it is not as the conserved centralspindlin complex. Surprisingly, *in vitro* and *in vivo* work show that the classically microtubule-associated Pavarotti binds directly to actin and has a non-canonical role directly regulating actin dynamics. We show that Pavarotti also works independently from Tumbleweed in several actin-related processes during the normal developmental process of oogenesis.

## Introduction

Centralspindlin is a conserved heterotetrameric complex composed of dimerized Pavarotti (Pav) and Tumbleweed (Tum) in Drosophila (CHO1/MKLP1 and MgcRacGAP in mammals, respectively) (D’Avino et al., 2015; Green et al., 2012; Mishima et al., 2002; Pollard and O’Shaughnessy, 2019; Somers and Saint, 2003; White and Glotzer, 2012). Pav is a member of a kinesin-like (kinesin-6) protein family of microtubule-dependent molecular motors, whereas Tum is a Rho GTPase activating protein (GAP) that exchanges the GTP-bound form of Rho family GTPase to the GDP-bound form (Adams et al., 1998; Crest et al., 2012; Goldstein et al., 2005; Somers and Saint, 2003; Sommi et al., 2010; Zavortink et al., 2005). The role of the centralspindlin complex in cytokinesis is highly conserved and well-studied in a wide variety of model organisms and cell lineages (Cheffings et al., 2016; D’Avino et al., 2015; Green et al., 2012; Pollard and O’Shaughnessy, 2019; White and Glotzer, 2012). In brief, during cytokinesis, centralspindlin organizes microtubules (MTs) in the central spindle region, associates with the Pebble (Pbl) Rho guanine nucleotide exchange factor (RhoGEF; Ect-2 in mammals), and moves along with Pbl to localize at the equator of the dividing cells (Basant and Glotzer, 2018; Cheffings et al., 2016; D’Avino et al., 2015; Green et al., 2012; Pollard and O’Shaughnessy, 2019; Somers and Saint, 2003; White and Glotzer, 2012). Subsequently, centralspindlin and Pbl activate Rho1 leading to formation of an actomyosin ring at the cortex of the cell equator (Basant and Glotzer, 2018; Cheffings et al., 2016; D’Avino et al., 2015; Green et al., 2012; Pollard and O’Shaughnessy, 2019; Somers and Saint, 2003; White and Glotzer, 2012). Tum, Pav, and Pbl are each essential for the spatial and temporal regulation of Rho1, and their recruitment to the equator is MT-dependent (Basant and Glotzer, 2018; Nishimura and Yonemura, 2006; Piekny et al., 2005; Somers and Saint, 2003; Yuce et al., 2005).

In addition to actomyosin ring formation, the centralspindlin complex also regulates MT dynamics during cytokinesis (D’Avino et al., 2015; Green et al., 2012; Pollard and O’Shaughnessy, 2019; White and Glotzer, 2012). In the early stages of cytokinesis, the centralspindlin complex localizes to the spindle midzone that is composed of anti-parallel bundled MTs. At later stages of cytokinesis, centralspindlin mediates the association of these bundled MTs in the spindle midzone with the plasma membrane to form the midbody structure that is important for completing the final stage of cytokinesis. Other examples of MT-dependent or -independent functions of centralspindlin include the migration of neurons in *C. elegans*, negative regulation of the WNT pathway, and MT sliding that promotes neurite outgrowth in Drosophila (Del Castillo et al., 2015; Falnikar et al., 2013; Jones et al., 2010). While in most contexts centralspindlin as a complex remains unchanged, a recent study in *C. elegans* showed that RhoGAP (Tum) independently regulates Rho1 activity in oocyte production by the syncytial germline (Lee et al., 2018).

Recently, we developed a model to study cell wound repair in the Drosophila syncytial embryo, in which the lateral side of nuclear cycle 4-6 Drosophila embryos are wounded and the repair process can be visualized in real-time by 4D confocal microscopy (Abreu-Blanco et al., 2011a; Abreu-Blanco et al., 2014; Nakamura et al., 2017). This model shares many features of cytokinesis, including an essential actomyosin contractile ring that is necessary for wounds to close (Abreu-Blanco et al., 2011a; Abreu-Blanco et al., 2014; Nakamura et al., 2018; Nakamura et al., 2017). To date, our work and the work of others has elucidated a cassette of molecular factors involved in cell wound repair that is strikingly similar to those used during cytokinesis (Cooper and McNeil, 2015; Dekraker et al., 2018; Nakamura et al., 2018; Sonnemann and Bement, 2012). In addition to the actomyosin ring, Rho1, Rac1, and Cdc42 are essential to wound healing where they exhibit distinct spatiotemporal patterns (Abreu-Blanco et al., 2014; Benink and Bement, 2005; Nakamura et al., 2017). Additionally, we recently found that Pbl (RhoGEF) and Tum (RhoGAP) are recruited to wounds and regulate the spatiotemporal dynamics of actin and myosin which form the actomyosin ring around the wound edge (Nakamura et al., 2017). Interestingly however, Pbl and Tum accumulate in distinct spatiotemporal patterns and Pbl regulates Cdc42 dynamics, rather than those of Rho1 as would be expected from cytokinesis studies. Additionally, Tum is required for the refinement of Rho1 and Rac1, but not Cdc42, suggesting that Pbl and the centralspindlin complex have separate roles in cell wound repair (Nakamura et al., 2017). Here, we investigate the role of Pav, the other member of the centralspindlin complex, in cell wound repair. We find that, in response to cell wounds, Pav is recruited in a distinct localization pattern and mutants exhibit a distinct phenotype compared with Tum and Pbl. Importantly, Pav localization at the wound is actin-dependent, can directly bind to actin, and functions during wound repair independent from the other centralspindlin complex members. Finally, we demonstrate that Pav regulation of actin dynamics independently of Tum is widespread.

## Results

### Pav, Tum, and Pbl are required, but have distinct roles, during cell wound repair

Recently, in elucidating the signaling pathways upstream of Rho family GTPase localization to the cell wound periphery, we uncovered two lines of evidence that suggested that the proteins forming the centralspindlin complex were involved in cell wound repair, but not as the canonical complex (Nakamura et al., 2017). Tum and Pbl did not exhibit the same localization patterns upon wounding and Tum and Pbl had distinct wound repair defects. However, as the core centralspindlin complex is composed of Tum and Pav, we were interested in if and how Pav was involved in cell wound repair.

Upon wounding, actin accumulates in a distinct pattern: 1) a highly-enriched actin ring bordering the wound edge, and 2) an elevated actin halo encircling the actin ring, which can be used a reference to orient other protein localizations (Fig. 1A) (Abreu-Blanco et al., 2011a; Abreu-Blanco et al., 2014; Nakamura et al., 2017). As we previously showed, Pbl accumulates in a diffuse ring-like array overlapping with the actin halo region, whereas Tum accumulates broadly around wounds overlapping with the actin-ring and halo (Fig. 1A-E; Video 1) (Nakamura et al., 2017). To examine the localization pattern of Pav, we wounded embryos expressing GFP-tagged Pav driven ubiquitously by the ubiquitin promoter (Minestrini et al., 2002) combined with a cherry-tagged actin reporter (sChMCA) (Abreu-Blanco et al., 2011a). Pav accumulates broadly around wounds overlapping with the actin ring and halo (Fig. 1A, F-G; Video 1). Both of Pav and Tum accumulate in the actin ring and halo, however, their localization patterns are different: Pav most strongly accumulates at the inner edge of actin ring, whereas Tum strongly accumulates in the actin halo region (Fig. 1A, D-G; Video 1).

**Fig 1.**
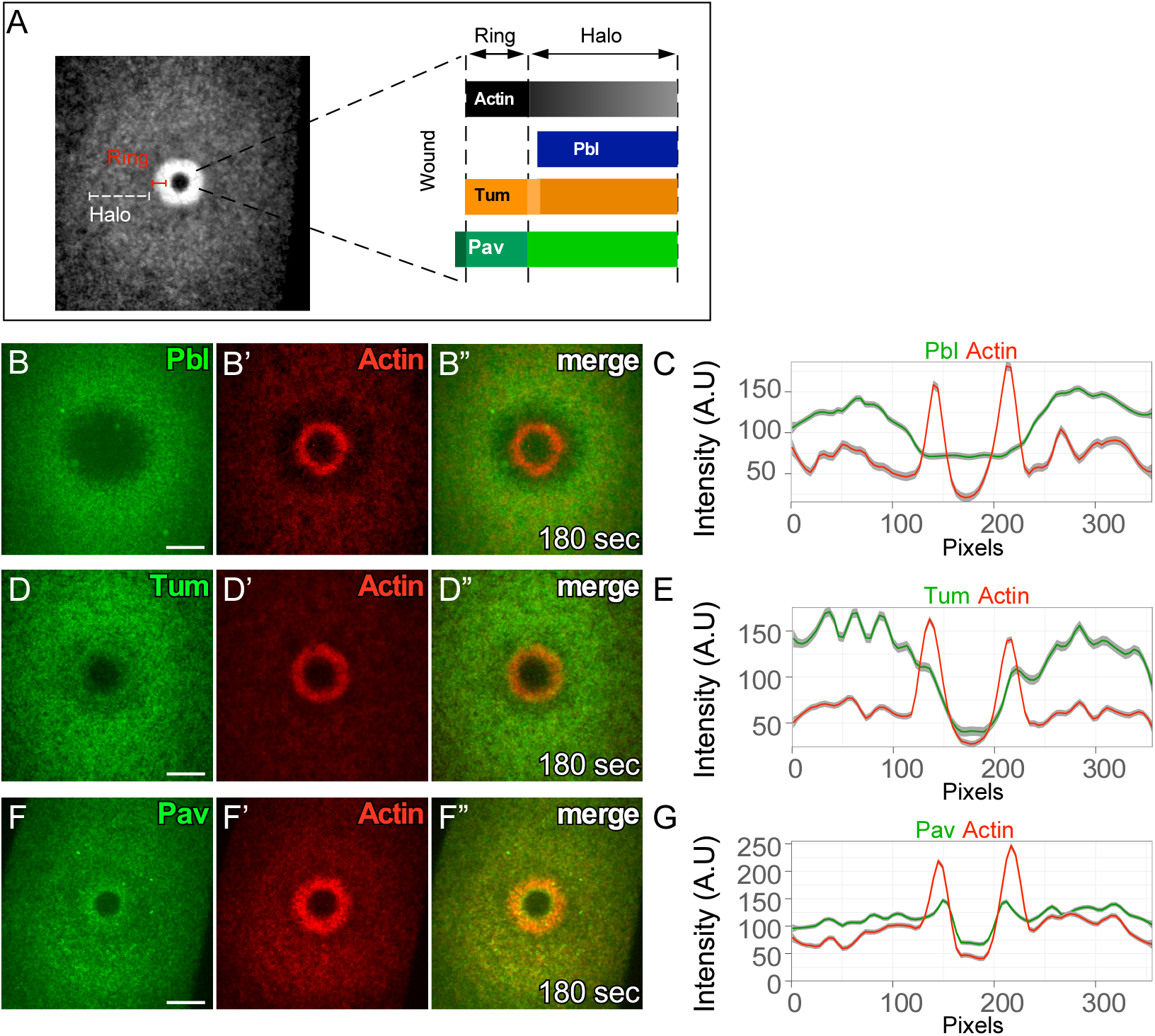
Pbl, Tum, and Pav exhibit distinct localization patterns in cell wound repair. (A) Confocal xy projection image from a laser wounded NC4-6 staged *Drosophila* embryo expressing an actin reporter (sGMCA). Schematic diagram summarizing the localization patterns of actin, Pbl, Tum, and Pav at the wound edge. (B-B’’) Confocal xy projection images from NC4-6 staged *Drosophila* embryo co-expressing an actin reporter (sChMCA) and Pbl-eGFP. (C) Fluorescence intensity (arbitrary units) profiles across the wound area in (B’’). (D-D’’) Confocal xy projection images from a NC4-6 staged *Drosophila* embryo co-expressing an actin reporter (sChMCA) and sfGFP-Tum. (E) Fluorescence intensity (arbitrary units) profiles across the wound area in (D’’). (F-F’’) Confocal xy projection images from a NC4-6 staged *Drosophila* embryo co-expressing an actin marker (sChMCA) and GFP-Pav. (G) Fluorescence intensity (arbitrary units) profiles across the wound area in (E). Scale bar: 20 μm. Time post-wounding is indicated.

We next were interested in comparing the wound repair phenotypes in *tum* and *pav* knockdown backgrounds using a fluorescent actin reporter (sGMCA or sChMCA) (Kiehart et al., 2000) that allows us to assay physical properties of the repair process (i.e., size, expansion, and closure rate) and defects in actin structures during this process. We generated *pav* knockdown embryos three different ways: 1) expressing two independent RNAi constructs for *pav* (HMJ02232 and GL01316) in the female germline using the GAL4-UAS system, 2) using the *wimp* mutation to generate reduced Pav expression in both the germline and soma (maternal contribution of Pav is reduced in trans-heterozygotes of *pav* and *wimp*, referred as reduced *pav*) (Parkhurst and Ish-Horowicz, 1991), and 3) expressing Pav^DEAD^, a previously described construct, which has a one amino acid substitution in the ATP binding site and overexpression of which causes dominant-negative phenotypes *in vivo* (Minestrini et al., 2003; Minestrini et al., 2002). Unfortunately, females with all combinations of maternal-GAL4s and RNAi lines do not produce any eggs. However, reduced *pav* and Pav^DEAD^ overexpressed by the nanos-GAL4 driver (referred to as Pav^DEAD^) females produce embryos that can be used to discern Pav function in wound repair. Both reduced *pav* and Pav^DEAD^ embryos exhibit similar phenotypes: delayed actin accumulation around the wound edge and delayed closure rate compared with control (*wimp*/+) embryos (Fig.2 A-C’, F-I; Video 2). In contrast to reduced *pav* mutants and Pav^DEAD^, *tum* mutant embryos obtained from RNAi knockdown/antibody injection (Nakamura et al., 2017) exhibit a different phenotype: wound overexpansion and a slightly increased closure rate (Fig. 2D-G; Video 2). Taken together, localization and mutant analyses indicate distinct roles for Pav and Tum.

**Fig 2.**
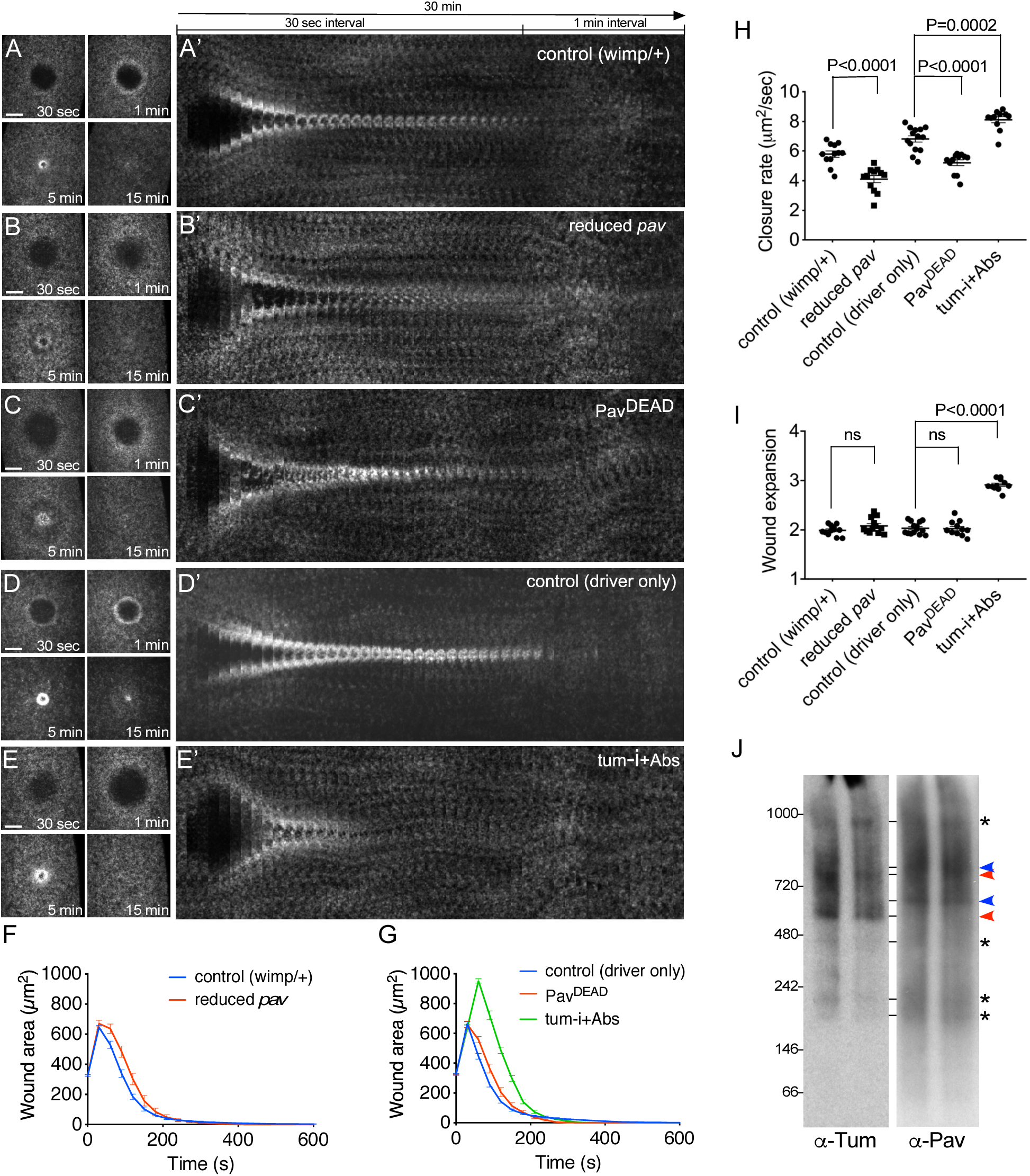
Pav and Tum mutants exhibit distinct phenotypes. (A-E). Actin dynamics (sGMCA or sChMCA) during cell wound repair in NC4-6 staged embryos: control (*wimp*/+) (A), reduced *pav* (*wimp* +/ + *pav*) (B), *pav^DEAD^* (C), control (GAL4 driver alone) (D), and *Tum-i*+Abs (E). (A’-E’) xy kymograph across the wound area depicted in (A-E), respectively. (F-G) Quantification of the wound area over time for control (*wimp*/+; n=12) (F), reduced *pav* (n=12) (F)*, pav^DEAD^* (n=12) (G), control (GAL4 driver alone; n=14) (G), and *Tum-i*+Abs (n=11) (G). (H-I) Quantification of wound closure speed (H) and wound expansion time (I) for control (*wimp*/+), reduced *pav, pav^DEAD^*, control (GAL4 driver alone), and *Tum-i*+Abs. Time post-wounding is indicated. Scale bar: 20 μm. Error bars represent ± SEM. Unpaired student’s t tests were performed in (H-I). (J) Western blot from BN-PAGE of Drosophila ovary lysates (the two lanes are loaded identically) probed with anti-Tum or anti-Pav antibodies (* indicates common Pav/Tum protein complexes; b lue arrowhead indicates Pav only protein complexes; red arrowhead indicates Tum only protein complexes).

To determine if these non-canonical Pav and Tum functions could be due to their functioning in separate complexes, we separated protein complexes from Drosophila ovary lysates using blue native-PAGE and probed the resulting western blots with anti-Pav or anti-Tum antibodies (Fig. 2J). While we observed multiple overlapping bands on both the Pav and Tum western blots (asterisks; Fig. 2J), several bands were unique to either Pav or Tum (blue or red arrowheads, respectively; Fig. 2J). Thus, in addition to acting as the centralspindlin complex, Pav and Tum can act in separate protein complexes.

### Rho family GTPases are recruited normally to wounds in *pav* mutant backgrounds

As centralspindlin complex is required to activate Rho1 during cytokinesis and spatiotemporal regulation of Rho family GTPases is indispensable for the formation of the actomyosin ring during cell wound repair, we examined if Pav regulates Rho family GTPase dynamics—with or without other RhoGEFs and RhoGAPs—in cell wound repair. We wounded embryos co-expressing a GFP-tagged actin reporter (sGMCA) and cherry-tagged Rho1, Rac1, or Cdc42 in control and reduced *pav* mutant backgrounds. In control embryos, Rho1 accumulates inside of the actin ring, whereas Rac1 and Cdc42 accumulate in the ring and halo regions (Fig. 3A-C’’; Video 3). These GTPases are still recruited to wounds and exhibit similar localization patterns to control embryos in the reduced *pav* mutant background, albeit the GTPases expression patterns are slightly different between control and reduced *pav* mutant backgrounds as a consequence of crosstalk between GTPase localization and the actin cytoskeleton, which is disrupted in this background (Fig. 3D-F’’; Video 3) (Abreu-Blanco et al., 2014; Nakamura et al., 2017). These results indicate that Pav is not required for the spatiotemporal Rho family GTPase localization during cell wound repair.

**Fig 3.**
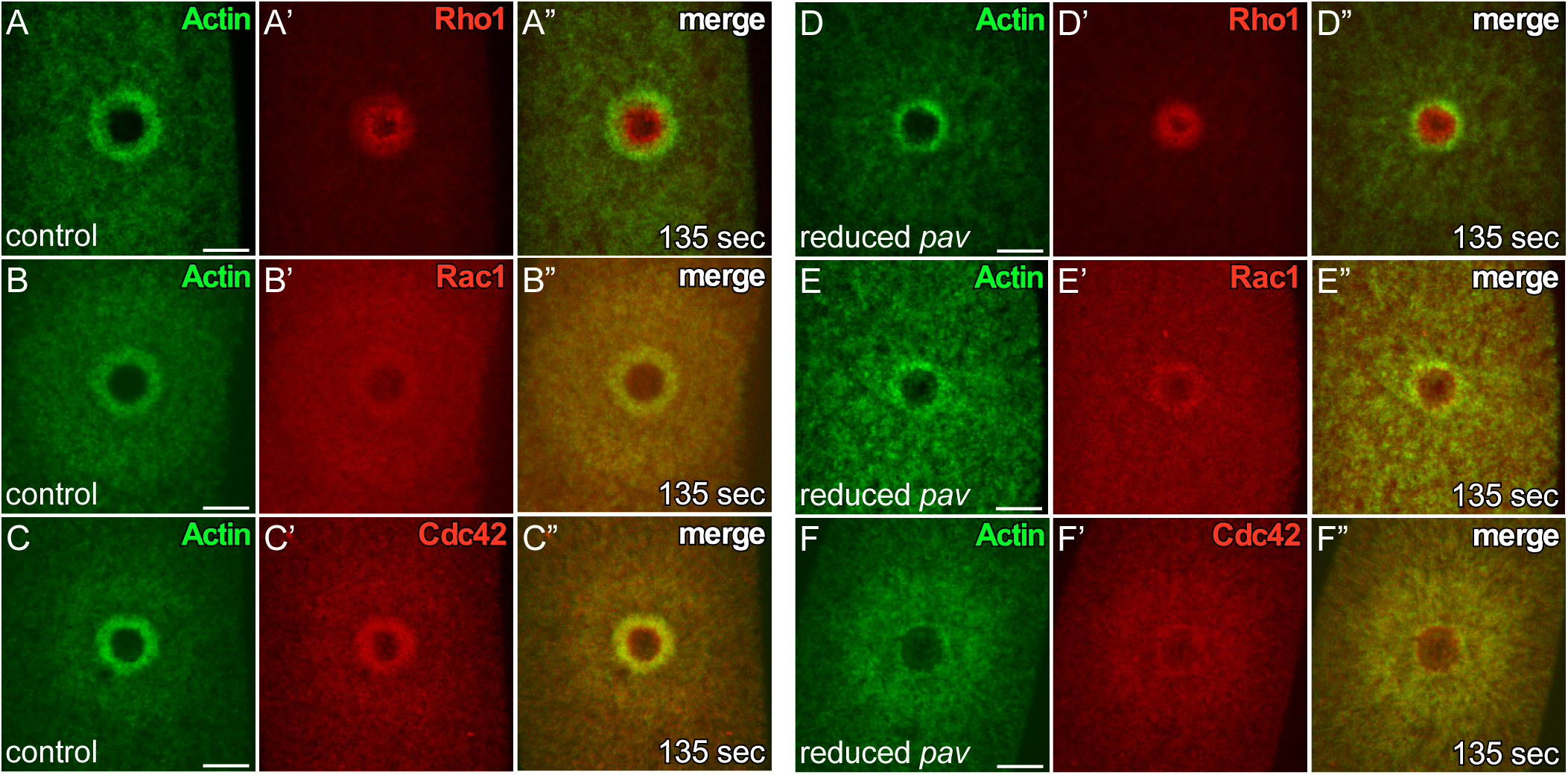
Spatial and temporal patterns of Rho family GTPases are not altered in reduced *pav* mutant backgrounds. (A-C’’) Localization of Rho family GTPases along with an actin reporter (sGMCA) in NC4-6 staged control embryos: ChFP-Rho1 (A-A’’), ChFP-Rac1 (B-B’’), and ChFP-Cdc42 (C-C’’). (D-F’’) Localization of Rho family GTPases along with an actin reporter (sGMCA) in NC4-6 staged reduced *pav* embryos: ChFP-Rho1 (D-D’’), ChFP-Rac1 (E-E’’), and ChFP-Cdc42 (F-F’’). Time post-wounding is indicated. Scale bar: 20 μm.

### Pav associates with the actin during cell wound repair

Kinesin-like proteins associate with MTs and are able to transport cargo proteins to where they are needed (Hirokawa et al., 2009; Lu and Gelfand, 2017; Vale, 2003), thus Pav may regulate the recruitment of actin regulators other than Rho family GTPases. Indeed, we previously found that injection of colchicine (inhibitor of MT polymerization) into embryos disrupts actin dynamics and wound closure (Abreu-Blanco et al., 2011a). To examine whether the kinesin motor function of Pav is required for actin dynamics, we wounded embryos co-expressing an actin reporter (sChMCA) and Pav-GFP upon injecting buffer, colchicine, or as a control, latrunculin B (LatB; inhibitor of actin polymerization). While Pav accumulation at the edge of the actin ring is disrupted in colchicine-injected embryos compared with buffer control, Pav is still recruited to wounds and its localization pattern is similar to that of actin (Fig. 4A-B’’; Video 4). In contrast to colchicine injection, we initially did not expect that actin disruption by LatB would affect Pav localization. Surprisingly, Pav localization is severely disrupted and largely mirrors the disrupted actin structures (Fig. 4C-D’’; Video 4). Since actin and MTs are often intimately associated with each other (cf. Dogterom & Koenderink, 2019), it is possible that MT disorganization by actin disruption could affect Pav localization. To test this possibility, we examined Pav localization upon injecting colchicine and LatB simultaneously, and in this case found that Pav still overlaps with the disrupted actin structures (Fig. 4E-F’’; Video 4). These results indicate that Pav associates with actin, rather than MTs, during cell wound repair.

**Fig 4.**
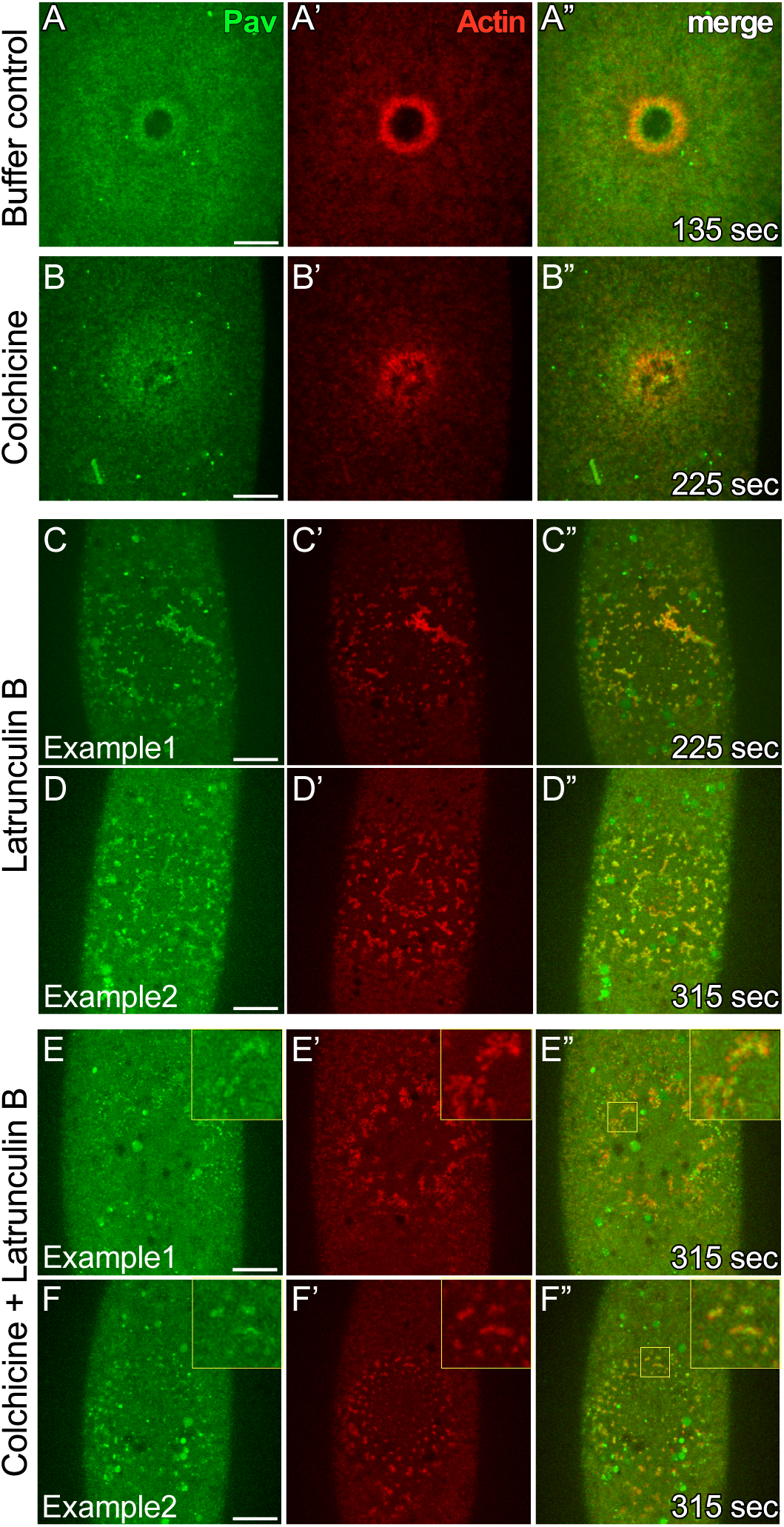
GFP-Pav overlaps with actin during cell wound repair. (A-F’’) Localization of GFP-Pav and an actin reporter (sChMCA) upon injecting buffer control (A-A’’), colchicine (B-B’’), LatB (C-D’’), or colchicine + LatB (E-F’’) in NC4-6 staged embryos. Time post-wounding is indicated. Scale bar: 20 μm.

### Pav binds, bundles, and crosslinks F-actin and MTs

Consistent with Pav associating with actin during cell wound repair, a previous study identified an actin binding site in the eighteenth exon of CHO1, one of two isoforms of MKLP-1, the mammalian homolog of Pav (Kuriyama et al., 2002). Although sequence alignment shows that this actin binding site is not conserved in Pav, our results suggest that Pav may bind directly to actin. To determine if Pav binds directly to F-actin, we first performed low-speed co-sedimentation of bacterially purified full-length Pav protein. Unfortunately, Pav protein goes to the pellet without F-actin and/or MTs present, perhaps due to its large size (>100kDa) and/or dimerization (data not shown). Hence, we next performed *in vitro* F-actin and MT bundling/crosslinking assays. *In vitro* polymerized F-actin or MTs distribute uniformly and are not bundled or crosslinked in the absence of additional factors (Fig. 5A-A’). Addition of bacterially-purified full-length Pav protein bundles MTs as reported previously (Tao et al., 2016) in the presence (Fig. 5B’-B”) or absence (Fig. 5D) of F-actin. Surprisingly, addition of Pav protein also bundles F-actin alone or in the presence of MTs (Fig. 5B-C), as well as crosslinks F-actin and MTs (Fig. 5B-B”). We examined the direct binding activity of Pav to actin and MTs by adding GFP-tagged Pav protein to actin and/or MTs bundled by Pav. GFP-tagged Pav binds directly to actin and MTs, alone (Fig. 5F-G”) and in the presence of each other (Fig. 5E-E””).

**Fig 5.**
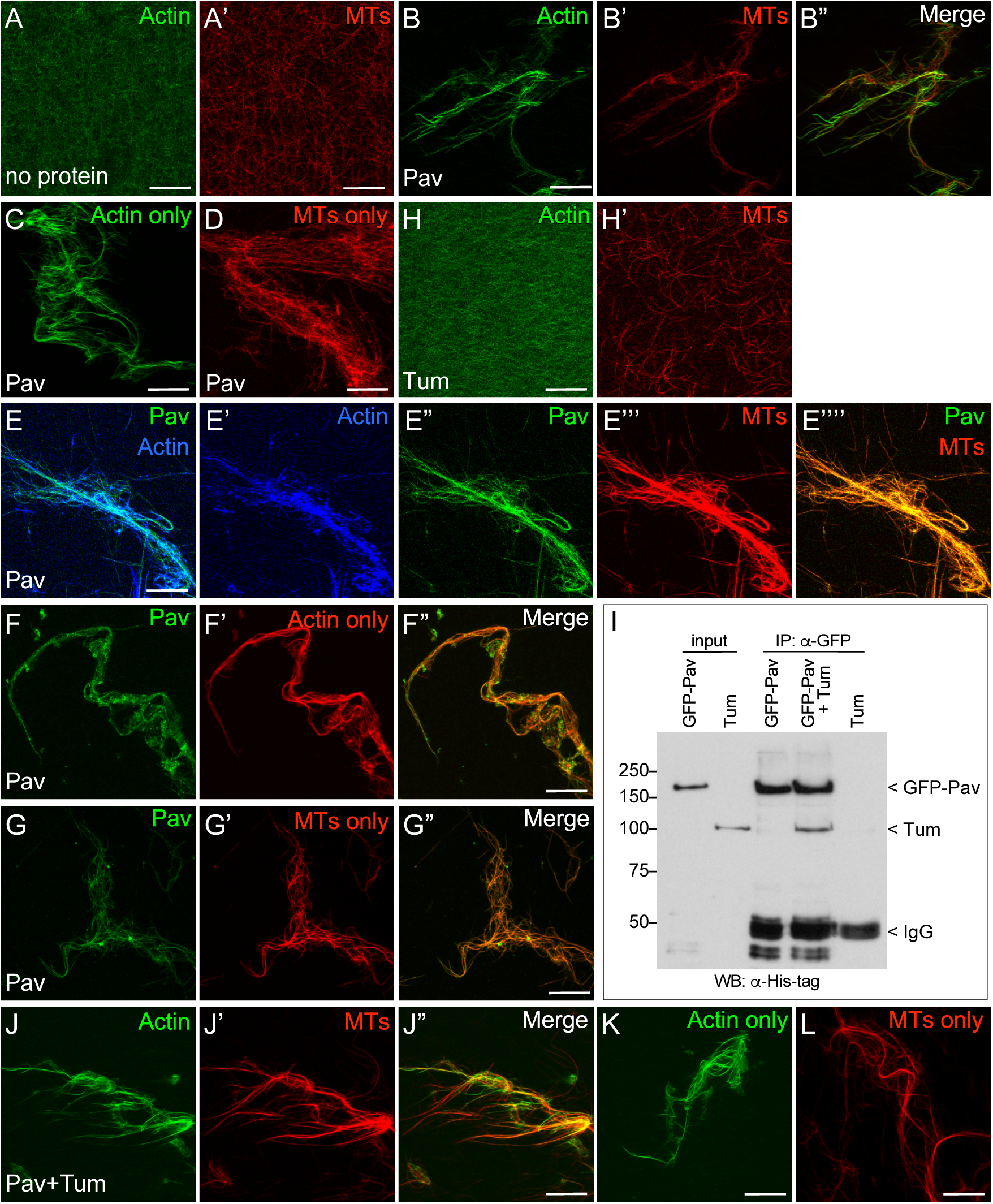
Pav bundles F-actin/MT and crosslinks them, even in the presence of Tum. (A-D) Stabilized actin and/or MTs were incubated with no protein (A-A’’) or full-length Pav protein (B-D). (E-G’’) Stabilized actin and/or MTs were bundled and cross-linked by full-length Pav protein then incubated with GFP-Pav. (H-K) Stabilized actin and/or MTs were incubated with Tum (H-H’) and Pav+Tum (1:2 molar ratio) (I-K). Final protein concentrations: Pav (100 nM), GFP-Pav (3 nM), and Tum (200 nM). (L) Western blots of co-immunoprecipitations from bacterially-purified His-tagged GFP-Pav and/or His-tagged Tum proteins with anti-GFP antibody. Anti-His antibody is used for detection of proteins. Scale bar: 30 μm.

Since previous studies had shown that Tum works as a molecular switch which activates the kinesin motor function of Pav (Davies et al., 2015; Tao et al., 2016; White et al., 2013), we examined whether the presence of Tum protein could affect the F-actin bundling and/or crosslinking activities of Pav. Bacterially-purified Tum protein alone does not bundle either F-actin or MTs (Fig. 5H-H’). We incubated Tum and GFP-tagged Pav proteins together to allow them to form the centralspindlin complex, and confirmed their physical association using co-immunoprecipitation (Fig. 5I). Allowing Tum to associate with Pav does not affect the bundling and/or crosslinking activities of Pav (Fig. 5J-L).

Overexpression of Pav^DEAD^ caused similar phenotypes to reduced *pav* mutants, so to further understand Pav’s bundling and crosslinking activity we purified Pav^DEAD^ and performed the same assays. Interestingly, Pav^DEAD^ protein cannot bundle actin/MTs or crosslink them (Fig. 6A-A’). To determine whether Pav^DEAD^ protein loses the ability to bundle actin because of an inability to bind actin, we added GFP-tagged Pav or Pav^DEAD^ protein to bundled and/or crosslinked actin and MTs. Importantly, the formin Cappuccino was used to bundle and/or crosslink actin and MTs so that the system was Pav-independent prior to its addition (Rosales-Nieves et al., 2006). Localization of GFP-tagged Pav^DEAD^ protein on actin/MTs, actin alone, and MTs alone (Fig. 6E-G”) is similar to GFP-tagged Pav (Fig. 6B-D”), suggesting that Pav^DEAD^ still can bind to actin and MTs. This loss of bundling phenotypes by Pav^DEAD^ is not due to a defect in dimerization as GST pull-down assays indicate that Pav^DEAD^ protein is still able to dimerize (Fig. 6H-I). Thus, Pav^DEAD^ protein loses actin/MT bundling activities, but retains its ability to bind to actin and MTs.

**Fig 6.**
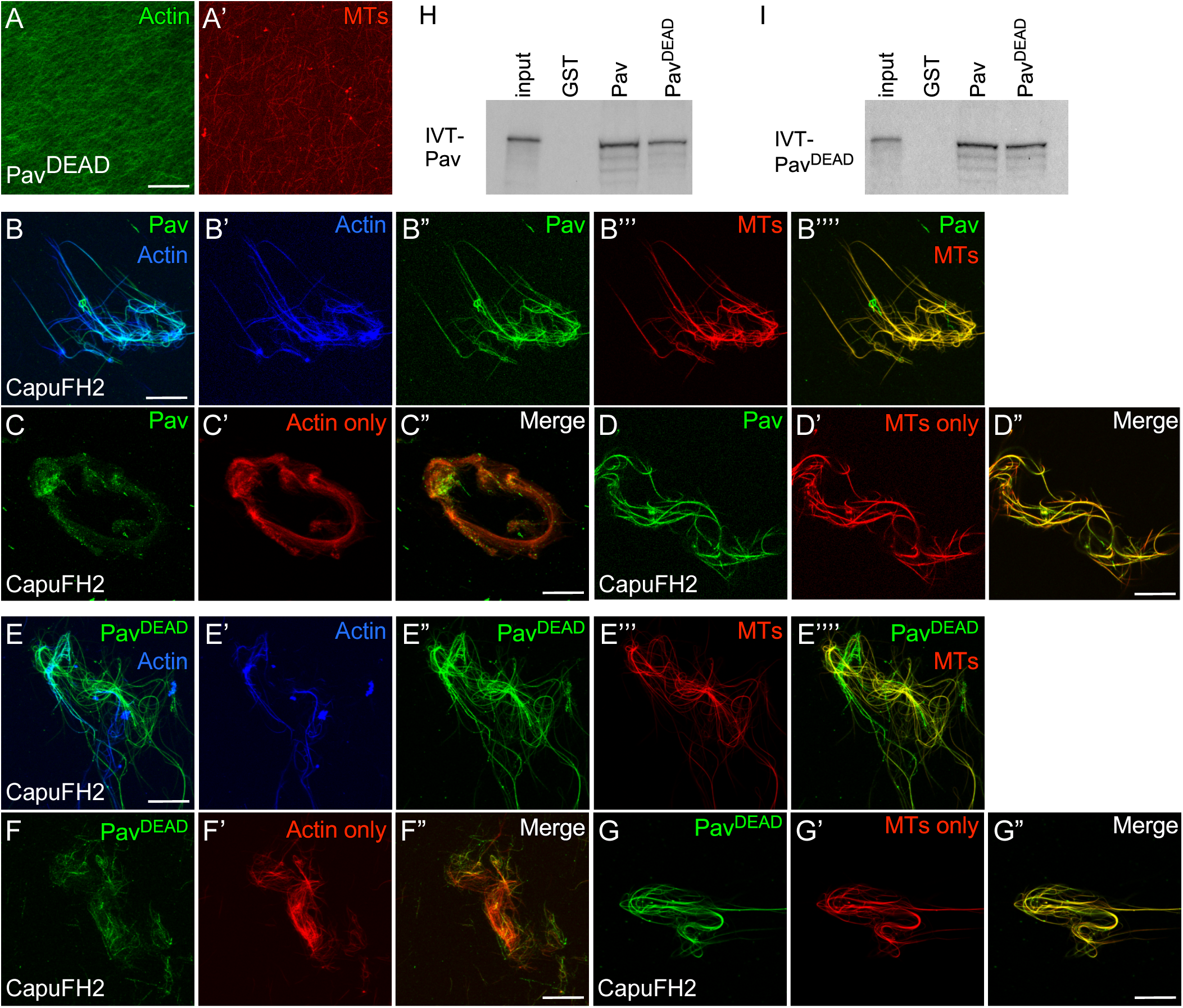
Pav ^DEAD^ protein cannot bundle and crosslink F-actin/MTs, but can bind to them. (A-A’) Stabilized actin and MTs were incubated with Pav ^DEAD^. (B-G’’) Stabilized actin and/or MTs were incubated with CapuFH2 to bundle actin and MTs and then added GFP-Pav (B-D’’) or GFP-PavDEAD (E-G’’). (H-I) GST pulldown assays with ^35^S-labeled *in vitro* translated Pav and Pav^DEAD^. The GST-Pav and Pav ^DEAD^ proteins were loaded. Final protein concentrations: Pav^DEAD^ (100 nM), CapuFH2 (500 nM), GFP-Pav (3 nM), and GFP-Pav ^DEAD^ (3 nM). Scale bar: 30 μm.

### Pav, but not Tum, accumulates at the oocyte cortex and reduced *pav* mutants exhibit premature ooplasmic streaming

We next were interested in exploring whether or not this actin-related (centralspindlin-independent) function of Pav might be required for other developmental processes. We had indirect evidence that this might be the case in oogenesis, as maternally driven RNAi lines produced embryos for Tum, but none in the case of Pav. Previous studies had also indicated that Pav localized to ring canals (Airoldi et al., 2011; Minestrini et al., 2002). These actin-rich structures, comprised of inner and outer actin rings, are created by incomplete cytokinesis in Drosophila oogenesis and function to connect the oocyte and nurse cells (Hudson and Cooley, 2002; Ong and Tan, 2010; Robinson et al., 1994; Warn et al., 1985) (Fig. 7A-J). To determine whether Pav in ring canals functions as the centralspindlin complex, we examined Tum and Pav localization in stage 7 egg chambers. We stained stage 7 egg chambers from flies expressing GFP-tagged Tum (driven by the ubiquitous sqh promoter) with antibodies to Pav and GFP (Nakamura et al., 2017). We found that while Tum and Pav are both present throughout the oocyte and nurse cells, they showed specific enriched localization at inner ring canals and in nurse cell nuclei (Fig. 7A-C’”). Interestingly, in addition to localizing to the inner ring canal, Pav also localizes with the outer actin rings (Fig. 7B-C’’’). We also observed Pav, but not Tum, enrichment at the oocyte cortex (Fig. 7A-A’’’, 7K-M”). It is not completely unexpected that both Tum and Pav localize at the ring canals as these were once sites of cytokinesis, but we were surprised that they showed only partially overlapping localizations. To follow up on this, we next looked at actin ring canal morphology in reduced *pav* and *tum* RNAi mutants. Based on their slightly different localizations, we suspected that these proteins were not entirely functioning together as the centralspindlin complex and would display only partially overlapping phenotypes. In comparison to WT ring canals, reduced *pav* mutants have a slightly disorganized inner actin ring with a broad, detached, and highly disorganized outer ring canal with actin spikes protruding into the cells (Fig. 7D-E, 7G-H, 7J). In contrast to this, *tum* RNAi mutants have a clearly distinct phenotype showing a mildly disorganized inner actin ring, prominent actin spikes into the nurse cells, and the F-actin filaments of the outer ring canal are detached from the cell cortex and are binding to the inner ring canal perpendicular to their normal orientation (Fig. 7F, 7I, 7J). Both protein localization and phenotypes of Tum and Pav suggests that the proteins are acting at least partially independently.

**Fig 7.**
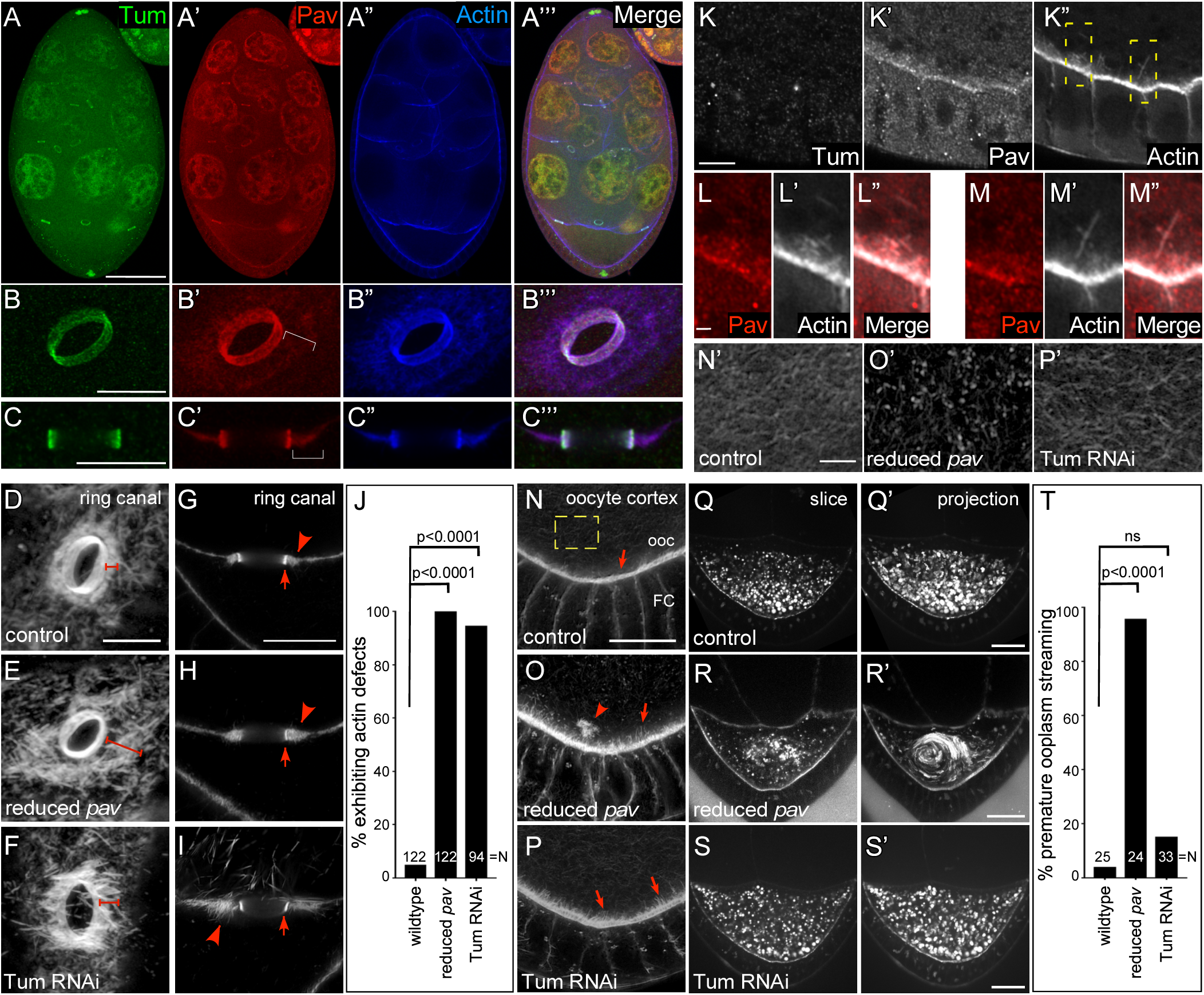
Actin-dependent Pav function is required for oogenesis. (A-A’’’) GFP-Tum expressing stage 7 egg chamber stained for Tum (anti-GFP), Pav (anti-Pav), and F-actin (phalloidin). (B-C’’’) Oblique view (B-B’’’) of the ring canal connecting two nurse cells and cross section (C-C’’’) of the ring canal connecting an oocyte and a nurse cell in the stage 7 egg chambers, stained for anti-Tum, anti-Pav, and F-actin. Brackets indicate outer actin ring canals. Scale bar: 5 µm. (D-F) Obliqu e view of the ring canal connecting two nurse cells in stage 7 egg chambers, stained for F-actin, in control (D), reduced *pav* (E), and Tum RNAi (F). (G-I) Cross-section of the ring canal connecting an oocyte and a nurse cell in stage 7 egg chamber, stained for F-actin, in control (G), reduced *pav* (H), and Tum RNAi (I). (J) Quantification of the percentage of egg chambers exhibiting actin defects. (K-K’’) Protein distribution at the GFP-Tum expressing oocyte cortex in the stage 7 egg chamber, stained for Tum (anti-GFP), Pav (anti-Pav), and F-actin (phalloidin). Scale bar: 5 µm. (L-M’’) High magnification views from two different areas in (K). Scale bar: 1 µm. (N-P) Posterior of stage 7 egg chambers, stained for F-actin, in control (N-N’), reduced *pav* (O-O’), and Tum RNAi (P-P’). Arrows indicate cortical actin projections from oocyte (ooc). Arrowhead indicates actin aggregation in the oocyte. (N’-P’) Cortical actin mesh structure in control (N’), reduced *pav* (O’), and Tum RNAi (P’) from (N-P). (Q-S’) Single time point and 30 min time point projections from time-lapse movies of stage 7 egg oocytes in control (Q-Q’), reduced *pav* (R-R’), and Tum RNAi (S-S’). (T) Quantification of the percentage of egg chambers exhibiting premature ooplasm ic streaming. N= for each genotype is indicated on bar plots and Fisher’s exact test were performed in (J) and (T); ns, not significant. Scale bars: 50 µm in (A-A’’’), 20µm in (Q-S’), 10 µm in (B-I, K-K’’, and N-P), and 1 µm in (L-M’’, N’-P’).

We were extremely interested in Pav’s role at the oocyte cortex as only Pav was enriched at this region and because we and others have previously shown that actin/MT bundling and crosslinking here are essential to prevent premature ooplasmic swirling (Dahlgaard et al., 2007; Liu et al., 2009; Lu et al., 2016; Rosales-Nieves et al., 2006; Wang and Riechmann, 2008). Since we observed that Pav binds F-actin filaments *in vitro* (Fig. 5), we utilized high resolution microscopy to determine that Pav also localizes to F-actin structures *in vivo* at the oocyte cortex (Fig. 7K-M”). In controls, cortical actin is highly organized with one layer of uniformly-sized of spike-like structures (Fig. 7K”, 7N). Reduced *pav* mutants exhibit disrupted cortical actin organization resulting in abnormally long actin filaments, actin aggregations within the ooplasm, and disrupted actin mesh (Fig. 7O-O’). Intriguingly, *tum* RNAi oocytes exhibited abnormal cortical actin, albeit less severely than reduced *pav*, with long actin filaments and separated cortex layers, despite no evidence of specific localization to this region (Fig. 7P). The actin mesh in these mutant oocytes was similar to that in wildtype (Fig. 7P’). This subtle phenotype may be indirect and due to the loss of Tum in other parts of the egg chamber.

We next looked in both mutant oocyte backgrounds to see if either Pav or Tum was required to prevent premature ooplasmic streaming (Fig. 7Q-T). Consistent with their disrupted cortical actin, ooplasmic actin aggregates, and disrupted ooplasm actin mesh, reduced *pav* mutants exhibit premature ooplasmic streaming (Fig. 7R-R’, 7T; Video 5). Interestingly, *tum* RNAi mutants do not exhibit premature ooplasmic streaming, consistent with retaining an ooplasm actin mesh similar to that in wildtype oocytes (Fig. 7Q-Q’, 7S-T; Video 5). Taken together with the ring canal findings, localization and mutant analyses indicate that Pav has Tum-independent, actin-related functions during normal oogenesis.

## Discussion

Pav is a kinesin-like protein that, along with the Tum RhoGAP protein, forms the centralspindlin complex that can bundle MTs and move along them during cytokinesis (D’Avino et al., 2015; Green et al., 2012; Pollard and O’Shaughnessy, 2019; White and Glotzer, 2012). Previously, CHO-1, the mammalian homolog of Pav, was shown to bind actin through a non-conserved protein domain, however the biological relevance of this activity is currently not known (Kuriyama et al., 2002). Here, we showed that Pav binds directly to F-actin and that actin-dependent, centralspindlin-independent functions of Pav are required for proper cell wound repair and oogenesis, which has important implications for both understanding the biological functions of Pav as well as the cell wound repair and oogenesis processes.

Cytoskeletal elements must be temporally and spatially coordinated for cells to carry out complex functions, including cell division and cell wound repair (Abreu-Blanco et al., 2011b; Agarwal and Zaidel-Bar, 2019; Basant and Glotzer, 2018; Bement and von Dassow, 2014; Cheffings et al., 2016; Chew et al., 2017; Chugh and Paluch, 2018; D’Avino et al., 2015; Dekraker et al., 2018; Dogterom and Koenderink, 2019; Green et al., 2012; Nakamura et al., 2018; Pollard and O’Shaughnessy, 2019; Sonnemann and Bement, 2012; Verboon and Parkhurst, 2015). MTs form a radial array pattern and associate with actin around wounds in *Xenopus* oocytes (Mandato and Bement, 2003), whereas MTs do not accumulate or become spatially arrayed in the Drosophila cell wound repair model (Abreu-Blanco et al., 2011a). Similarly, while the accumulation of actin and myosin at the wound periphery is regulated by Rho family GTPases and required for proper cell wound closure, the means by which they assemble into an actomyosin ring and translocate to close wounds varies between cell wound models (Abreu-Blanco et al., 2014; Burkel et al., 2012). *Xenopus* oocytes have been proposed to employ an actin treadmilling mechanism wherein actin is continually polymerized at the interior of the actomyosin ring and depolymerized at the outer edge of the actomyosin ring to promote ring translocation and wound closure (Burkel et al., 2012). In contrast, the actomyosin ring at the wound periphery in Drosophila syncytial embryos is sensitive to myosin inhibitors and functions as a contractile array (Abreu-Blanco et al., 2014). We find that Pav localizes at edge of the wound, overlapping with the inner edge of actomyosin ring. Previously, we found that Rho1 GTPase and the formin Diaphanous (Dia) similarly accumulate inside of the actin ring during cell wound repair, where they are both required for actomyosin ring assembly. Since Dia can polymerize F-actin, newly polymerized actin formed inside of the actin ring would need to be aligned and assembled into the actomyosin ring. Pav could be one of proteins that is required for actin alignment, assembly, actin bundling, actin/MT crosslinking, and/or actomyosin ring assembly during this wound repair process. In addition to organizing actin structures, Pav might be involved in maintaining uniform tension around the actomyosin ring or in the generation of contractile forces by sliding bundled actin filaments as non-muscle myosin does. Indeed, MKLP1 has been shown to organize anti-parallel bundled MTs during cell division and then slide these MTs past each other to generate force in the spindle midzone (Mastronarde et al., 1993; Nislow et al., 1992; Sharp et al., 1996).

In addition to cell wound repair, we find that localization patterns and mutant phenotypes of Pav and Tum in egg chambers are not identical. Ring canals, the stable intracellular bridges between germ cells, are composed of two highly organized actin-rich structures—the inner and outer ring canals (Hudson and Cooley, 2002; Ong and Tan, 2010; Robinson et al., 1994; Warn et al., 1985). As ring canals are the result of incomplete cytokinesis, we expected that Pav and Tum would function together as the centralspindlin complex at these structures. We were surprised to find that while both Pav and Tum were enriched along with actin at inner ring canals, only Pav was enriched with actin at outer ring canals. The inner ring canal defects we observe might be due to disruption of cytokinesis during the early stages of oogenesis dependent on the roles of Pav and Tum as the centralspindlin complex, whereas Pav’s centralspindlin-independent F-actin binding, bundling, and/or crosslinking activities are needed for proper organization of the outer ring canals. Pav may be required to organize and compact actin in the outer ring canals since reduced *pav* mutants exhibit detached and disorganized actin, similar to that observed in *wash* mutants that has also been shown to bundle and crosslink actin/MTs (Liu et al., 2009; Verboon et al., 2018). In contrast to Pav, the aberrant attachment of the outer ring canal to the inner ring canal in *tum* mutants is likely indirect since Tum is not enriched at the outer ring canals. One possibility is that Rho1 and/or Rac1 activities are disrupted in *tum* mutants since we have shown that Tum is necessary to refine Rho1 and Rac1 patterns during cell wound repair (Nakamura et al., 2017). Interestingly, a recent study found that each protein of the centralspindlin complex in *C. elegans* has independent roles during germline development: both CYK-4 (RacGAP) and ZEN-4 (kinesin-6) localize to intercellular bridges that are formed by incomplete cytokinesis, whereas CYK-4, but not ZEN-4, is essential for germline development (Lee et al., 2018).

We show that Pav organizes another type of actin structure during oogenesis: the oocyte cortex. Interestingly, Rho1 GTPase, as well as the branched actin nucleation factor Wash and the linear actin nucleation factors Capu (formin) and Spire, are similarly enriched at the oocyte cortex, have been shown to regulate actin and MT organization/crosstalk at the oocyte cortex, and, when mutant, result in premature ooplasmic streaming in stage 7-8 oocytes (Liu et al., 2009; Rosales-Nieves et al., 2006; Verboon et al., 2018). It is striking, but unclear, why so many different actin regulatory proteins are required non-redundantly to regulate actin and/or MT dynamics at the oocyte cortex thereby preventing premature ooplasmic streaming. One possibility is that Pav has a major function in stabilizing the F-actin network by bundling actin or crosslinking F-actin/MTs that have already been polymerized by Wash, Capu, or Spire, since a primary function of these proteins is to nucleate F-actin filaments. Consistent with this, we observed detached actin aggregates within the ooplasm in reduced *pav* mutants. Another possibility is that Pav might regulate Rho1 activity in this context, thereby disrupting downstream effectors. In contrast to Pav, our results suggest that Tum is not essential for preventing premature ooplasmic streaming, albeit *tum* mutants exhibit mild actin defects at the oocyte cortex. Premature ooplasmic streaming has been proposed to occur when an actin mesh present in stage 7 oocytes is disrupted such that MTs are free to move past one another (Dahlgaard et al., 2007; Lu et al., 2016; Wang and Riechmann, 2008). Consistent with this model, the actin mesh in reduced *pav* mutant oocytes is disrupted, whereas the actin mesh in *tum* mutant oocytes is indistinguishable from that of wild type. Since Tum is not enriched at the oocyte cortex, the mild cortical actin disruption we observe might be an indirect effect.

In summary, we show an unexpected actin-dependent function for the kinesin-like protein Pav during cell wound repair and oogenesis, suggesting that Pav can change modes between primarily actin- or primarily MT-dependent functions *in vivo*. We have shown that Tum presence is not the switch that distinguishes between these two modes of action. As we find that Pav and Tum form both overlapping and independent complexes, identification of other complex members is a future priority in order to understand the molecular basis of this switch, as well as how widespread these non-canonical functions are.

## Materials and Methods

### Fly stocks and genetics

Flies are cultured and crossed at 25ºC on yeast-cornmeal-molasses-malt medium. All fly stocks were treated with tetracycline, then tested by PCR to ensure that they did not harbor Wolbachia. The following stocks were used: OregonR (wild type), sGMCA (Kiehart et al., 2000), sChMCA (Bloomington #35519, #35520, #35521; (Abreu-Blanco et al., 2011a), ChFP-Rho1 (Bloomington #52280, #52281, #52282; (Abreu-Blanco et al., 2014), ChFP-Rac1 (Bloomington #76266, #76267; (Abreu-Blanco et al., 2014), ChFP-Cdc42 (Bloomington #42236, #42237; (Abreu-Blanco et al., 2014), sqh-sfGFP-Tum (Bloomington #76264, #76265; (Nakamura et al., 2017), sqh-Pbl-eGFP (Bloomington #76257, #76258; (Nakamura et al., 2017), Ubi-GFP-Pav (Minestrini et al., 2002), UASp-GFP-Pav^DEAD^ (Minestrini et al., 2002), UAS-RacGAP50C-dsRNA (Bloomington #6439; (Nakamura et al., 2017), TRiP.HMJ02232 (Pav RNAi, Bloomington #42573), TRiP.GL01316 (Pav RNAi, Bloomington #43963), pav^963^ (Bloomington #23926), and RpII140^wimp^ (Bloomington #5874; (Parkhurst and Ish-Horowicz, 1991).

To knockdown genes, RNAi lines were driven maternally using the GAL4-UAS system with P{matalpha4-GAL-VP16}V37 for driving Tum RNAi (Bloomington #7063) or P{w[+mC]=GAL4::VP16-nos.UTR}MVD1 for driving Pav RNAi or UASp-Pav^DEAD^ (Bloomington #4937). Reduced *pav* embryos were obtained from trans-heterozygous females by crossing *pav^963^* females to *RpL140^wimp^* males. All RNAi experiments were performed at least twice from independent genetic crosses and ≥10 embryos were examined unless otherwise noted.

### Embryo handling and preparation

Nuclear cycle (NC) 4-6 Drosophila embryos were collected from 0-30 min at room temperature (22ºC). Embryos were hand dechorionated, placed onto No. 1.5 coverslips coated with glue, and covered with Series 700 halocarbon oil (Halocarbon Products Corp).

### Laser wounding

All wounds were generated with a pulsed nitrogen N2 Micropoint laser (Andor Technology Ltd., Concord, MA, USA) tuned to 435 nm and focused on the cortical surface of the embryo. A region of interest was selected in the lateral midsection of the embryo and ablation was controlled by MetaMorph. On average, ablation time was less than 3s, and time-lapse imaging was initiated immediately. Occasionally, a faint grid pattern of fluorescent dots is visible at the center of wounds that arises from damaged vitelline membrane that covers embryos.

### Drug and antibody injections

Pharmacological inhibitors and antibodies were injected from the dorsal side into the center of NC4-6 staged Drosophila embryos, and laser wounding was performed 5 min post-injection. The following inhibitors were used: Latrunculin B (0.5 mM; EMD); colchicine (25 mM; Sigma-Aldrich). Colchicine was prepared in injection buffer (5 mM KCl, 0.1 mM NaP pH6.8). Latrunculin B was prepared in injection buffer + 10% DMSO. Injection buffer + 10% DMSO is used as control. The following antibodies were obtained from the Developmental Studies Hybridoma Bank: anti-Tum (1H5) and anti-Tum (2B6). All antibodies were dialyzed in PBS and concentrated prior to injection. The two Tum Abs were mixed and injected into embryos at 120 ng/µl.

### Microscopy

All imaging was performed at room temperature (22ºC). The following microscopes were used:

1) For live imaging: Revolution WD systems (Andor Technology Ltd., Concord, MA, USA) mounted on a Leica DMi8 (Leica Microsystems Inc., Buffalo Grove, IL, USA) with a 63x/1.4 NA objective lens and controlled by MetaMorph software. Images and videos were acquired with 488 nm and 561 nm, using an Andor iXon Ultra 897 or 888 EMCCD cameras (Andor Technology Ltd., Concord, MA, USA). All images for cell wound repair were 17-20 µm stacks/0.25 µm steps. For single color, images were acquired every 30 sec for 15 min and then every 60 sec for 25 min. For dual green and red colors, images were acquired every 45 sec for 30-40 min. Live imaging for premature ooplasm streaming was performed as previously described (Verboon et al., 2018).

2) For bundling/crosslinking assays and fixed tissues: Zeiss LSM 780 spectral confocal microscope (Carl Zeiss Microscopy GmbH, Jena, Germany) fitted with Zeiss 20x/0.8 Plan-Apochromat, 40x/1.3, and 63×/1.4 oil Plan-Apochromat objectives. FITC (Alexa 488) fluorescence was excited with the 488 nm line of an Argon laser, and detection was between 500-580 nm. Red (Alexa 568) fluorescence was excited with the 561 nm line of a DPSS laser and detection was between 580-642 nm. Far-red (Phalloidin 633) fluorescence was excited with the 633 line of an Argon laser, and detection was between 643-723 nm. Pinhole was typically set to 1.0 Airy Units. Confocal sections were acquired at 0.25-1.0 micron spacing. Super-resolution images were acquired using an Airyscan detector in Super Resolution mode and captured confocal images were then processed using the Airyscan Processing feature on the Zen software provided by the manufacturer (Carl Zeiss Microscopy GmbH, Jena, Germany).

### Image processing, analysis, and quantification

All images were analyzed with Fiji (Schindelin et al., 2012). Measurements of wound area were done manually. To generate xy kymographs, all time-lapse xy images were cropped to 5.8 µm x 94.9 µm and then each cropped image was lined up. To generate fluorescent profile plots by R, 10 pixel sections across the wound were generated using Fiji as we showed previously (Nakamura et al., 2017). Line profiles from the left to right correspond to the top to bottom of the images unless otherwise noted.

Quantification of the wound expansion and closure rate was performed as follows: wound expansion was calculated with max wound area/initial wound size. Closure rate was calculated with two time points, one is tmax that is the time of reaching maximum wound area, the other is t<half that is the time of reaching less than the half of maximum wound area. Using these time points, average speed was calculated with (wound area at tmax – wound area at t<half)/tmax-t<half. Generation of all graphs, student’s t test, and Fisher’s exact test were performed with Prism 7.0a (GraphPad Software Inc.).

### Protein expression

GFP-Pav, Pav, GFP-Pav^DEAD^, and Pav^DEAD^ cDNAs were amplified as 5’SalI-3’NotI fragments from Ubi-GFP-Pav or UASp-GFP-Pav^DEAD^ flies and then cloned into a double tag pGEX vector (GST and His; (Liu et al., 2009). Tum cDNA was amplified as a 5’BamHI-3’NotI from a sqh-sfGFP-Tum construct and then cloned into a double tag pGEX vector. Protein expression were performed as previously described (Rosales-Nieves et al., 2006). CapuFH2 protein purification was performed as previously described (Rosales-Nieves et al., 2006). For other protein purification, cells were lysed by sonication in T300G5 buffer (50 mM Tris pH 7.6, 300 mM NaCl, 5%glycerol, 1mM DTT, 1% Triton X-100) with 50 mM imidazole and Complete protease inhibitor tablets (Roche). Lysates were centrifuged at 10,000 g for 30 min and the supernatants were coupled to Fastflow nickel-sepharose (GE) for 3 hours at 4ºC. The matrix was washed three times with T300G5 buffer with 50 mM imidazole, eluted by T300G5 with 1 M imidazole and flash frozen.

### Blue-native Polyacrylamide Gel Electrophoresis (BN-PAGE)

BN-PAGE was performed using a Novex Native Page Bis-Tris Gel System (Invitrogen) following the manufacturer’s protocol. Ovary lysates were spun at 16,100 x g for 10 min (4°C) followed by spinning through Ultrafree MC HV centrifugal filters (Millipore) at 11,000 x g for 5 min (4°C). The filtered ovary lysate was then mixed with 4X Native PAGE Sample Buffer (Invitrogen), loaded onto 3-12% Bis-Tris Native PAGE gels, and electrophoresed using 1X Native PAGE Running buffer system (Invitrogen). For clarity, two identically loaded lanes of lysate are shown. NativeMark Protein standard (Invitrogen) was used as the molecular weight marker. Lysate samples were transferred to PVDF membrane and blotted according to standard procedures. The following primary antibodies were used: anti-Tumbleweed (1:10 each of mouse monoclonal 1H5 and 2B6; Developmental Studies Hybridoma Bank, Iowa City, IA) and anti-Pavarotti (1:3000 of rabbit polyclonal; (Adams et al., 1998). Secondary antibodies used were: donkey-anti-mouse-horseradish peroxidase (1:15000; Jackson ImmunoResearch Laboratories) and donkey-anti-rabbit-horseradish peroxidase (1:30,000; Jackson ImmunoResearch Laboratories). Antibodies were visualized using an ECL Kit (ThermoScientifc).

### GST pull-down assays

GST pull-down assays were performed as previously described (Rosales-Nieves et al., 2006). In vitro translated (IVT-) proteins were synthesized using a TNT quick-coupled transcription-translation kit (Promega, Madison, WI). For synthesizing the Pav and Pav^DEAD^ proteins, each ORF was cloned into a pCite4 vector (MilliporeSigma, Burlington, MA) using standard cloning techniques. In each case, 10% input is shown. All experiments were performed at least two times from individually synthesized proteins.

### F-actin/microtubule bundling and crosslinking assays

F-actin and MT bundling/crosslinking assays were performed as previously described (Rosales-Nieves et al., 2006). All experiments were performed at least three times using independently purified proteins.

### Immunoprecipitations

GFP-Pav and Tum proteins were incubated with mouse anti-GFP antibody (Roche) overnight at 4°C. Protein G sepharose (15 µl) was then added in 0.5 ml Carol Buffer (50 mM Hepes pH7.9, 250 mM NaCl, 10 mM EDTA, 1 mM DTT, 10% glycerol, 0.1% Triton X-100) + 0.5 mg/ml BSA + protease inhibitors (Complete EDTA-free Protease Inhibitor cocktail; Roche) and the reaction allowed to proceed for 2 hours at 4°C. The beads were washed one time with Carol Buffer + BSA and two times with Carol Buffer alone. Analysis was conducted using SDS-PAGE followed by Western blots. Antibodies used for the IP western blots are as follows: anti-His monoclonal (1:3000 dilution; ThermoFisher Scientific) and goat anti-mouse HRP (1:15,000 dilution; Jackson ImmunoResearch Laboratories Inc.). All IP experiments were performed at least three times.

### Immunostaining of ovaries

Female flies were fattened on yeast for 2 days and then ovaries were dissected and fixed as described above. After 3 washes with PBS plus 0.1% Triton X-100, ovaries were permeabilized in PBS plus 1% Triton X-100 at room temperature for 2 h. Ovaries were washed 3 times with PAT [1× PBS, 0.1% Tween-20, 1% bovine serum albumin (BSA), 0.05% azide] then blocked in PAT at 4°C for 2 h. Antibodies were used at the following concentrations: mouse anti-GFP monoclonal (1:100; Roche), rabbit anti-Pav polyclonal (1:250; provided by Dr. David Glover, (Adams et al., 1998)), and the ovaries incubated for 48 hours at 4ºC. Ovaries were washed three times with XNS (1× PBS, 0.1% Tween-20, 0.1% BSA, 4% normal goat serum) for 40 min each, then incubated with Alexa Fluor 488-conjugated secondary antibodies and Alexa Fluor 568-conjugated secondary antibodies (1:1000; Invitrogen) overnight at 4°C. Ovaries were washed with PTW (1× PBS, 0.1% Tween-20), incubated with Alexa Fluor 633-conjugated Phalloidin at 0.005 units/μl (Molecular Probes/ Invitrogen, Rockford, IL) at room temperature for 1 h, and then washed with PTW. Ovaries were dissected into individual ovarioles, then mounted on slides in Slowfade Gold (Invitrogen). A minimum of two biological replicates were performed for each condition.

### Actin visualization in ovaries

Female flies were fattened on yeast for 2 days and then ovaries were dissected into cold PBS. Ovaries were fixed using 1:6 fix/heptane for 10 min. Fix is: 16.7 mM KPO4 pH 6.8, 75 mM KCl, 25 mM NaCl, 3.3 mM MgCl2, 6% formaldehyde. Ovaries were washed 3 times with PBS plus 0.1% Triton X-100, and then incubated in PBS plus 0.5% Triton X-100 and Alexa Fluor 568-conjugated Phalloidin at 0.005 units/μl (Molecular Probes/ Invitrogen, Rockford, IL) at room temperature for 1 h. Ovaries were washed with PTW (1× PBS, 0.1% Tween-20) 10 times for 10 min each, then dissected into individual ovarioles and mounted on slides in Slowfade Gold with DAPI medium (Invitrogen, Rockford, IL). A minimum of two biological replicates were performed for each condition.

## Acknowledgements

We thank David Glover, Helen McNeill, Yonit Tsatskis, Bina Sugumar, Parkhurst lab members, the Bloomington Stock Center, the Harvard Transgenic RNAi Project, and the Developmental Studies Hybridoma Bank for advice, antibodies, DNAs, flies, and other reagents used in this study.

## Author Contributions

All authors performed experiments. MN, JMV, and SMP contributed to the design and interpretation of the experiments, and to the writing of the manuscript.

## Author ORCIDs

Mitsutoshi Nakamura – http://orcid.org/0000-0001-7879-3176

Jeffrey M. Verboon – http://orcid.org/0000-0002-4454-6043

Clara L. Prentiss – http://orcid.org/0000-0002-8877-0119

Susan M. Parkhurst – http://orcid.org/0000-0001-5806-9930

## VIDEO LEGENDS

**Video 1. Pav and Tum exhibit distinct localization patterns in cell wound repair.**

(A-B) Time-lapse confocal xy images from Drosophila NC4-6 staged embryos co-expressing an actin reporter (sChMCA, red) along with sfGFP-Tum (green) (A) or GFP-Pav (green) (B). Time post-wounding is indicated. UW: unwounded.

**Video 2. Pav and Tum mutants exhibit distinct phenotypes.**

(A-E) Time-lapse confocal xy images from Drosophila NC4-6 staged embryos expressing an actin marker (sGMCA or sChMCA): (A) control (wimp/+), (B) control (Gal4 driver/+), (C) reduced *pav*, (D) Pav^DEAD^, and (E) Tum RNAi + Abs. Time post-wounding is indicated. UW: unwounded.

**Video 3. Rho family GTPases are recruited to cell wounds in reduced *pav* mutants.**

(A-F) Time-lapse confocal xy images from Drosophila NC4-6 staged embryos co-expressing an actin reporter (sGMCA, green) along with Rho family GTPase in control (A-C) and reduced *pav* (D-F): Ch-Rho1 (A and D, red), Ch-Rac1 (B and E, red), and Ch-Cdc42 (C and F, red). Time post-wounding is indicated. UW: unwounded.

**Video 4. GFP-Pav overlaps with actin during cell wound repair.**

(A-F) Time-lapse confocal xy images from Drosophila NC4-6 staged embryos co-expressing an actin reporter (sChMCA, red) with GFP-Pav (Green) upon injecting buffer (A), colchicine (B), latrunculin B (C-D), or colchicine + latrunculin B (E-F). Time post-wounding is indicated. UW: unwounded.

**Video 5. Reduced *pav*, but not *tum*, mutants exhibit premature ooplasmic streaming during oogenesis.**

(A-C) Time-lapse movies of stage 7 oocytes shown in Figure 7: control (A), reduced *pav* (B), and Tum RNAi (C). Time is indicated.

